# Acute joint inflammation induces a sharp increase in the number of synovial fluid EVs and modifies their phospholipid profile

**DOI:** 10.1101/2023.05.05.539599

**Authors:** Laura Varela, Chris H.A. van de Lest, Janneke Boere, Sten F.W.M. Libregts, Estefania Lozano–Andres, P. René van Weeren, Marca H.M. Wauben

**Affiliations:** Division Equine Sciences, Department of Clinical Sciences, Faculty of Veterinary Medicine, Utrecht University, Yalelaan 1, Utrecht, 3584 CM, The Netherlands; Division Cell Biology, Metabolism & Cancer, Department of Biomolecular Health Sciences, Faculty of Veterinary Medicine, Utrecht University, Utrecht, 3584 CM, The Netherlands; Department of Orthopaedics, University Medical Center Utrecht, Heidelberglaan 100, Utrecht, 3584 CX, The Netherlands; Division of Infectious Diseases & Immunology, Department Biomolecular Health Sciences, Faculty of Veterinary Medicine, Utrecht University, Yalelaan 1, Utrecht, 3584 CL, The Netherlands

**Author notes:** Corresponding author; *E-mail address:* (M.H.M.Wauben).

**Keywords:** Extracellular vesicles, Lipidomics, Synovial fluid, Equine, Inflammation, Synovitis

## Abstract

Inflammation is the hallmark of most joint disorders. However, the precise regulation of induction, perpetuation, and resolution of joint inflammation is not entirely understood. Since extracellular vesicles (EVs) are critical for intercellular communication, we aim to unveil their role in these processes. Here, we investigated the EVs’ dynamics and phospholipidome profile from synovial fluid (SF) of healthy equine joints and from horses with lipopolysaccharide (LPS)–induced synovitis. LPS injection triggered a sharp increase of SF–EVs at 5–8hr post–injection, which started to decline at 24h post–injection. Importantly, we identified significant changes in the lipid profile of SF–EVs after synovitis induction. Compared to healthy joint–derived SF–EVs (0h), SF–EVs collected at 5, 24, and 48h post–LPS injection were strongly increased in hexosylceramides. At the same time, phosphatidylserine, phosphatidylcholine, and sphingomyelin were decreased in SF–EVs at 5h and 24h post–LPS injection. Based on the lipid changes during acute inflammation, we composed specific lipid profiles associated with healthy and inflammatory state–derived SF–EVs. The sharp increase in SF–EVs during acute synovitis and the correlation of specific lipids with either healthy or inflamed states–derived SF–EVs are findings of potential interest for unveiling the role of SF–EVs in joint inflammation, as well as for the identification of EV–biomarkers of joint inflammation.

## 1. Introduction

The ability to move and exercise is paramount to the well–being of humans and animals. Therefore, disorders of the musculoskeletal system have a huge impact on the quality of life and, in human society, form an important economic burden in terms of health care costs [1]. Arthritis is one of the most common types of musculoskeletal ailments, with as most important forms being rheumatoid arthritis (RA) and osteoarthritis (OA). The incidence of the latter is increasing sharply in the rapidly aging populations of many societies [2].

Inflammation is now commonly accepted as an essential hallmark of all clinically relevant joint disorders [3]. Inflammation is a response from the organism designed to cope with external stimuli, such as infection or tissue injury. Its prominence varies among joint diseases based on the underlying cause of arthritis, with RA having a more prominent inflammatory character than OA. Osteoarthritis is characterized by a different form of inflammation with intermittent inflammatory bouts that are pivotal in the maintenance of the vicious cycle of tissue degeneration leading to diminishing tissular resistance with higher vulnerability to further damage [4].

Extracellular vesicles (EVs) are cell–derived bilipid membrane vesicles that are shed from the plasma membrane or secreted from multivesicular bodies in the extracellular environment. They play a crucial role in intercellular communication and regulation of inflammation by the delivery of bioactive cargoes such as proteins, RNAs, and bioactive lipids that can affect the function of targeted cells [5–9]. Since EVs are present in substantial amounts in the synovial fluid (SF) of patients with RA and OA, the interest to decipher their potential role in joint pathology, as well as to exploit EVs as biomarkers for these diseases or as therapeutics for joint repair and regeneration, has increased considerably [10–13].

Until now, EV biology has been almost exclusively focused on the identification of the molecular composition of EV–associated RNAs and proteins in an attempt to determine their cell of origin, the message they convey, and their target cells to ultimately unveil their (patho)physiological function or to define EV–biomarkers for disease or EV–based therapeutics. By viewing it only as the envelope of the EV content, current research has so far largely neglected the dynamics of the EV lipid bilayer membrane by viewing it only as the envelope of the EV content. Nevertheless, lipids are crucial structural elements in charge of EV biogenesis, secretion, and function [14], and changes in the cellular environment can induce changes in the phospholipidome of EVs [15]. The EV membrane comprises glycerophospholipids (GPLs) and sphingolipids (SLs). In addition, Evs transport lipid–related molecules, e.g., PLA2 enzymes and lipid mediators, and carry lipid second messengers, e.g., phosphatidic acid and ceramides, connected to EV biogenesis [16].

Hence, lipids can be ligands that trigger signal transduction pathways, intermediates of signaling cascades, or substrates for lipid phosphatases and kinases [17–19]. Thus, a detailed understanding of the lipidic composition of SF–derived EVs can shed light on the role of EVs in joint diseases and might fuel further research into the development of potential diagnostic and therapeutic applications. To address this question, we used a well–described model for the induction of an acute, yet transient, synovitis in horses [20–22]. We herewith aim to study the effect of acute synovitis on both the dynamics of EVs and, more in detail, on the lipid composition and changes in the phospholipid profile of SF–derived EVs.

## 2. MATERIALS AND METHODS

### 2.1. Ethical Considerations

In vivo experiments were in compliance with the Dutch Experiments on Animals Act and conform to the European Directive 2010/63/EU. Furthermore, they were authorized by the Utrecht University Animal Ethics Committee (DEC), the Animal Welfare Body (IvD), and the Central Authority for Scientific Procedures on Animals (CCD) under license number 10800.

### 2.2. Experimental LPS–induced acute synovitis in adult horses

#### In vivo study 1

A transient and fully reversible acute synovitis was induced in n=6 adult Warmblood mares by aseptic intra–articular injection into the left or right middle carpal joint of 0.5 ng LPS from E. Coli O55:B5 (Sigma Aldrich, St. Louis, MO, USA) [21]. Synovial fluid (SF) was collected at 0 hours pre–LPS–injection, at 8h, 24h, and 168h post–LPS–injection, and stored at –80°C. Samples of in vivo study 1 were used for Fig. 1E.

**Figure 1:**
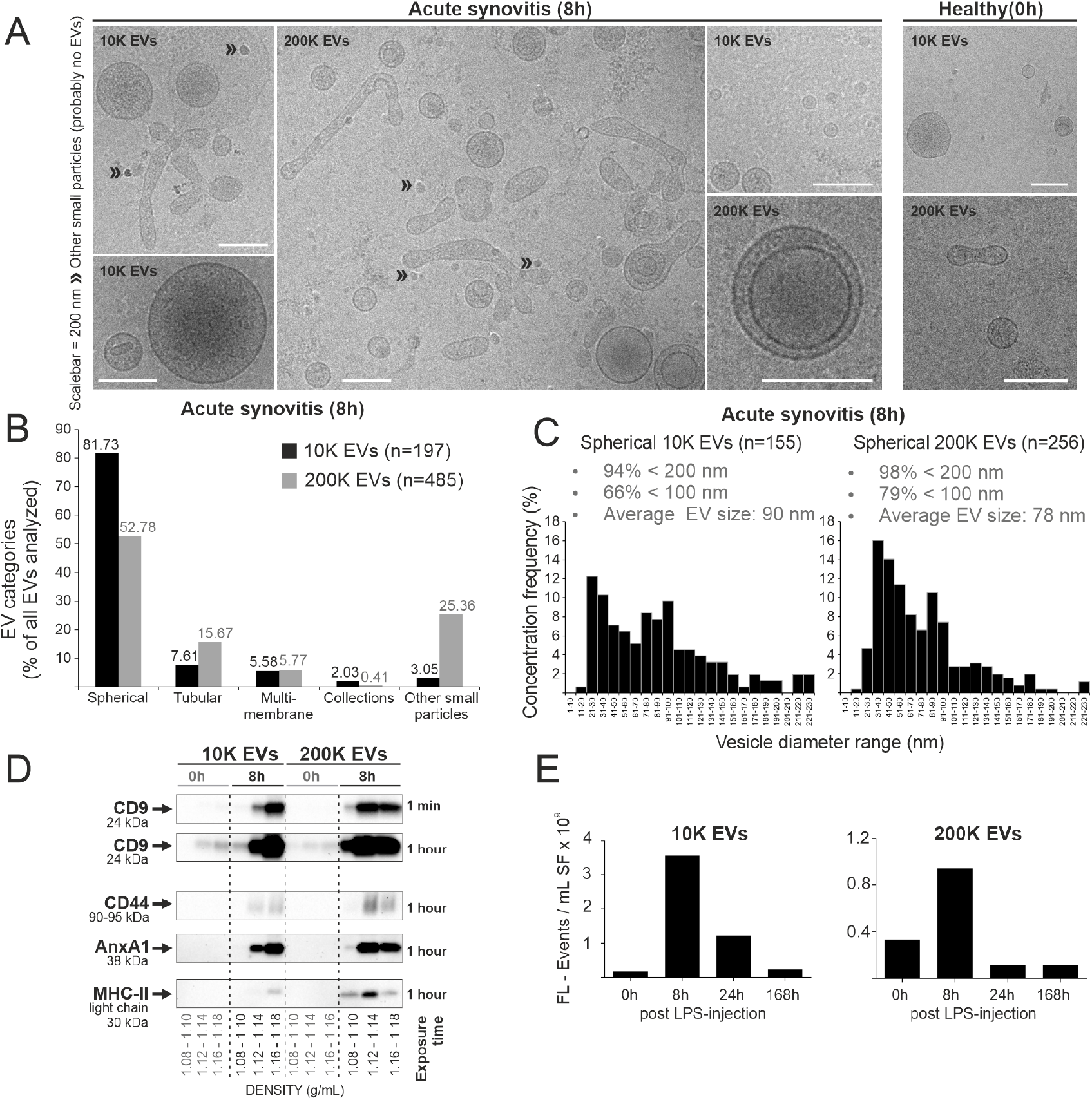
Qualitative and quantitative analysis of EVs isolated from joints with acute synovitis or healthy joints. **A)** Representative cryo–TEM images of 10K and 200K 8h SF–EVs, compared to the healthy situation analyzed identically as previously performed by Boere [24]. **B)** Quantification of typical EV morphologies observed by cryo–TEM in 8h SF–EVs. The 200K centrifugation step also retrieved a significant number of particles without a clear membrane, referred to as “other small particles”; examples are indicated with » in 1A (these particles are not considered EVs; possibly these are lipoproteins (Sodár et al., 2016). **C)** Cryo–TEM defined size distribution of spherical 10K and 200K SF– derived EVs at 8h (post–LPS injection). **D)** Western blot detection of CD9, CD44, AnxA1, and MHC–II of EV–containing sucrose gradient fractions **E)** EV concentration in SF calculated as the sum of single fluorescent events measurements (PKH67+ events) by flow cytometry in EV–containing sucrose gradient densities (1.10 to 1.20 g/mL). Bars are pools of 3 animals (n=1). For each time point, 0.2 mL cell–free SF per animal was pooled to acquire a starting volume of 0.6 mL SF per condition for the isolation of EVs. FL – Events: Fluorescent Events.

#### In vivo study 2

Synovitis was induced in n=8 adult Warmblood horses (two mares, six geldings) by injecting 0.25 ng LPS from E. Coli O55:B5 into the left and right middle carpal and tarsocrural joints [22]. Synovial fluid was collected at 0 hours pre–LPS–injection, at 8h and 24h post–LPS–injection, and stored at –80°C. Samples of in vivo study 2 were used for Fig. 1A–C.

#### In vivo study 3

Synovitis was induced in one adult trotter mare by a single injection of 0.7 ng LPS from E. Coli O55:B5 into the left middle carpal joint and the right tarsocrural joint. Synovial fluid was collected at 0 hours pre–LPS–injection and 8 hours post–LPS–injection and stored at –80°C. Samples of in vivo study 3 were used for Fig. 1D and Suppl. Fig. 2.

#### In vivo study 4

Synovitis was induced in n=8 female skeletally mature Warmblood horses by injecting into the right middle carpal joints 2.5 ng LPS from E. Coli O55:B5 [23].

Synovial fluid was collected at 0 hours pre–LPS injection and 5, 24, and 48 hours post–LPS–injection. Samples of in vivo study 4 were used for Fig. 2–5 and Suppl. Fig. 3–6. Additionally, synovial fluid of n=3 adult warmblood horses was collected post–mortem. These patients were donated to science by their owners at the Utrecht University Equine Hospital and were euthanized for reasons unrelated to joint diseases. Synovial fluid samples of these patients were stored at –80°C and used for Fig. 2–5 and Suppl. Fig. 4–6.

**Figure 2:**
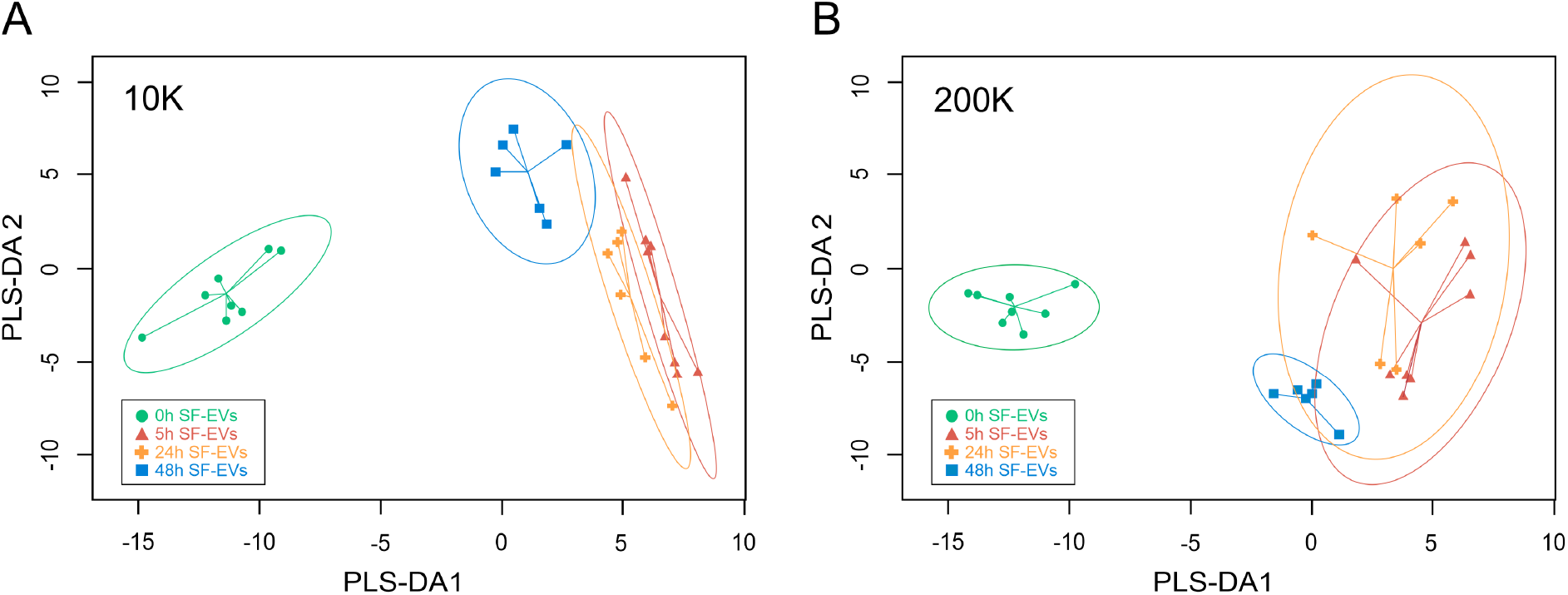
Partial least square discriminant analysis (PLS–DA) of lipids isolated from synovial fluid–derived EVs from healthy joints (baseline, 0 hours) and LPS–induced synovitis at 5 hours, 24 hours, and 48 hours. **A)** PLS–DA 1 vs. PLS–DA 2 plot demonstrating evident discrimination of healthy vs. post–LPS–injection EV samples. Each dot represents one EV–containing density fraction, of which each animal included contributed with two fractions. **A)** Lipid profile of 10k EVs. EV–containing density fractions 1.16 g/mL and 1.14 g/mL were used for the analysis. **B)** Lipid profile of 200k EVs. EV–containing density fractions 1.14 g/mL and 1.12 g/mL were used for the analysis. From each animal, both 10k and 200k EVs were isolated. Lipids were obtained from EVs isolated by differential ultracentrifugation and floated in sucrose density gradients. Healthy samples pre–LPS–0h (green circle; n=4 horses), synovitis samples post–LPS–5h (red triangle; n=4 horses), synovitis–resolution samples post–LPS–24h (orange cross; n=3 horses), resolution samples post–LPS–48h (blue square; n=3 horses). The ellipses around the dots indicate the 95% confidence intervals.

**Figure 3:**
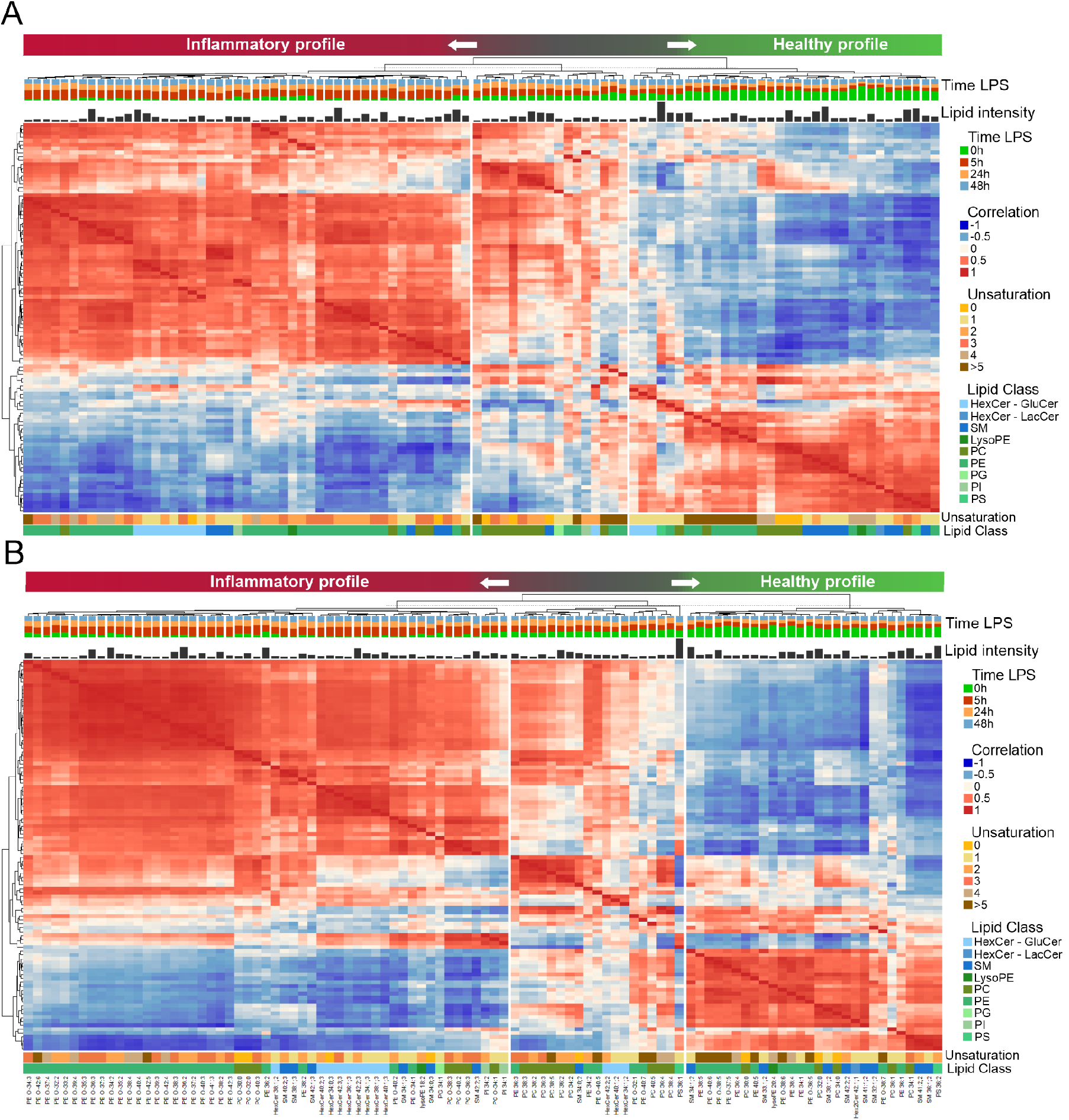
Lipid species correlation of synovial fluid–derived EVs. Combined heatmap (cluster dendrogram) of Lipid–Lipid Spearman correlations between the 100 most abundant lipid species in all EV sample groups. Lipid order is based on hierarchical clustering. On top of the figure is our proposed interpretation of the clustering. Below are the cluster dendrogram, the group distribution (upon which we based the proposed interpretation), the relative lipid intensity of each species, and the heatmap. Under the heatmap, the degree of saturation, the lipid class of each lipid, and the respective annotation of each lipid species are indicated. Lipids were obtained from SF–EVs pelleted with consecutive 10,000xg (10K) and 200,000xg (200K) ultracentrifugation steps followed by purification with sucrose density gradients. Healthy samples pre–LPS–0h (n=4), synovitis samples post–LPS–5h (n=4), synovitis–resolution samples post–LPS–24h (n=3), resolution samples post–LPS–48h (n=3). **A)** Heatmap of 10,000xg centrifuged EV. Density fractions 1.16 and 1.14 g/mL of each EV sample were used for the analysis. **B)** Heatmap of 200,000xg centrifuged EV. Density fractions 1.14 – 1.12 g/mL of each EV sample were used for the analysis.

**Figure 4:**
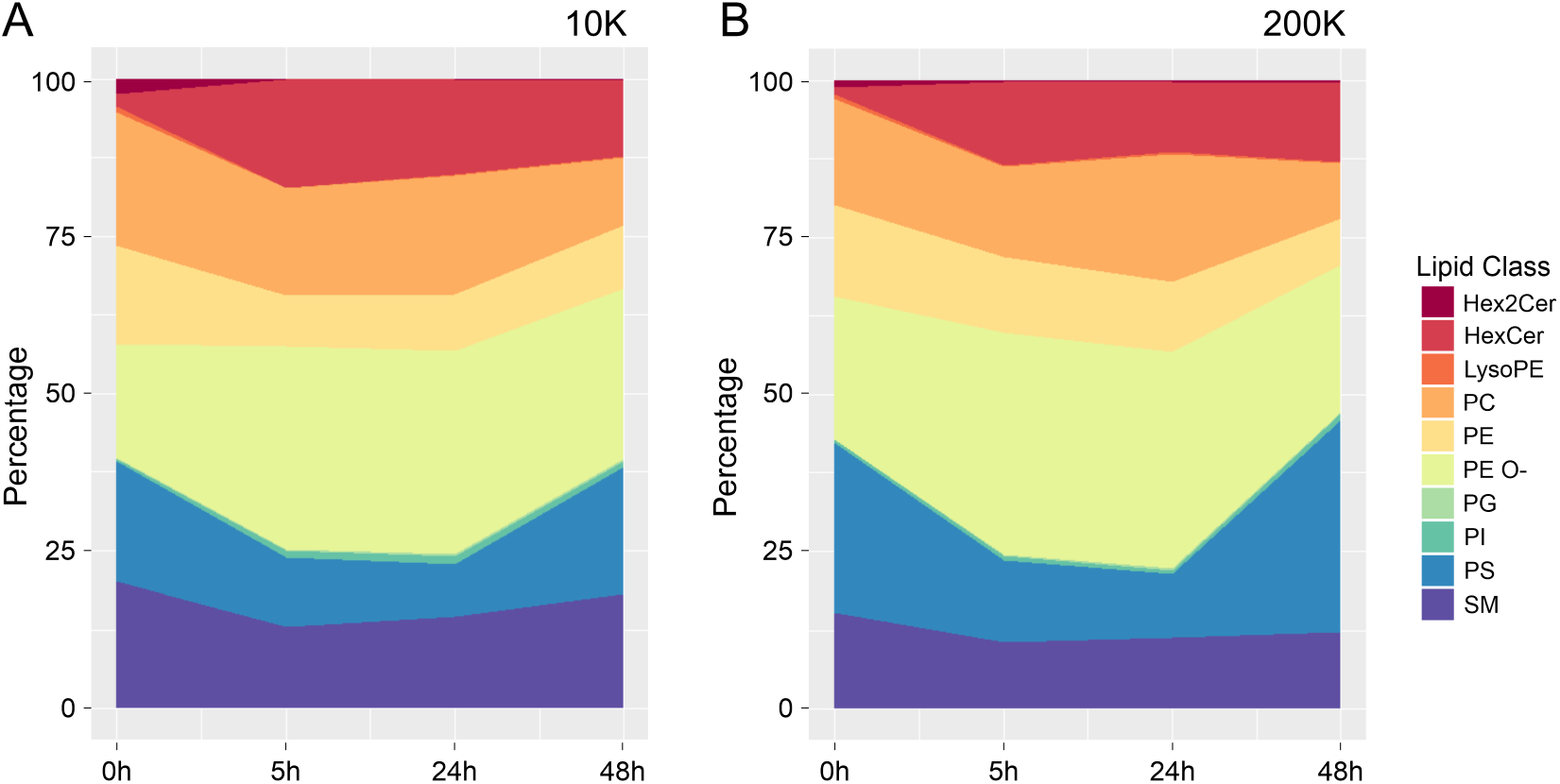
Changes in EV lipid classes during inflammation. Fractional composition plot of synovial fluid–derived EVs at 0 hours (pre–LPS), 5 hours, 24 hours, and 48 hours (post–LPS). The y–axis is the percentage (%) fraction of each lipid class; the x–axis represents the four time points of SF harvesting. Lipids were obtained from SF–EVs centrifugated at 10,000xg (10K) and 200,000xg (200K), followed by flotation of EVs in sucrose density gradients. Healthy samples were taken pre–LPS–0h (n=4), synovitis samples at post–LPS–5h (n=4), synovitis–resolution samples at post–LPS–24h (n=3), and resolution samples at post–LPS–48h (n=3). All samples are composed of two fractions. All fractions/samples were averaged within the four time points. Abbreviations: Hex2Cer (di–hexosylceramide); HexCer (hexosylceramide); LysoPE, (lysophosphatidylethanolamine); PC, (phosphatidylcholine); PE, (ester–linked phosphatidylethanolamine); PE O–, (ether–linked phosphatidylethanolamine); PG, (phosphatidylglycerol); PI, (phosphatidylinositol); PS, (phosphatidylserine); SM, (sphingomyelin). **A)** Fractional composition plot of 10K EVs. EV– containing density fractions 1.16 and 1.14 g/mL of all donors per time–point were averaged for this analysis. **B)** Fractional composition plot of 200k EVs. EV–containing density fractions 1.14 and 1.12 g/mL of all donors per time–point were averaged for this analysis. Noteworthy, if ceramides were detected during analysis, these were discarded during lipid annotation because their retention time was too close to the solvent front.

**Figure 5:**
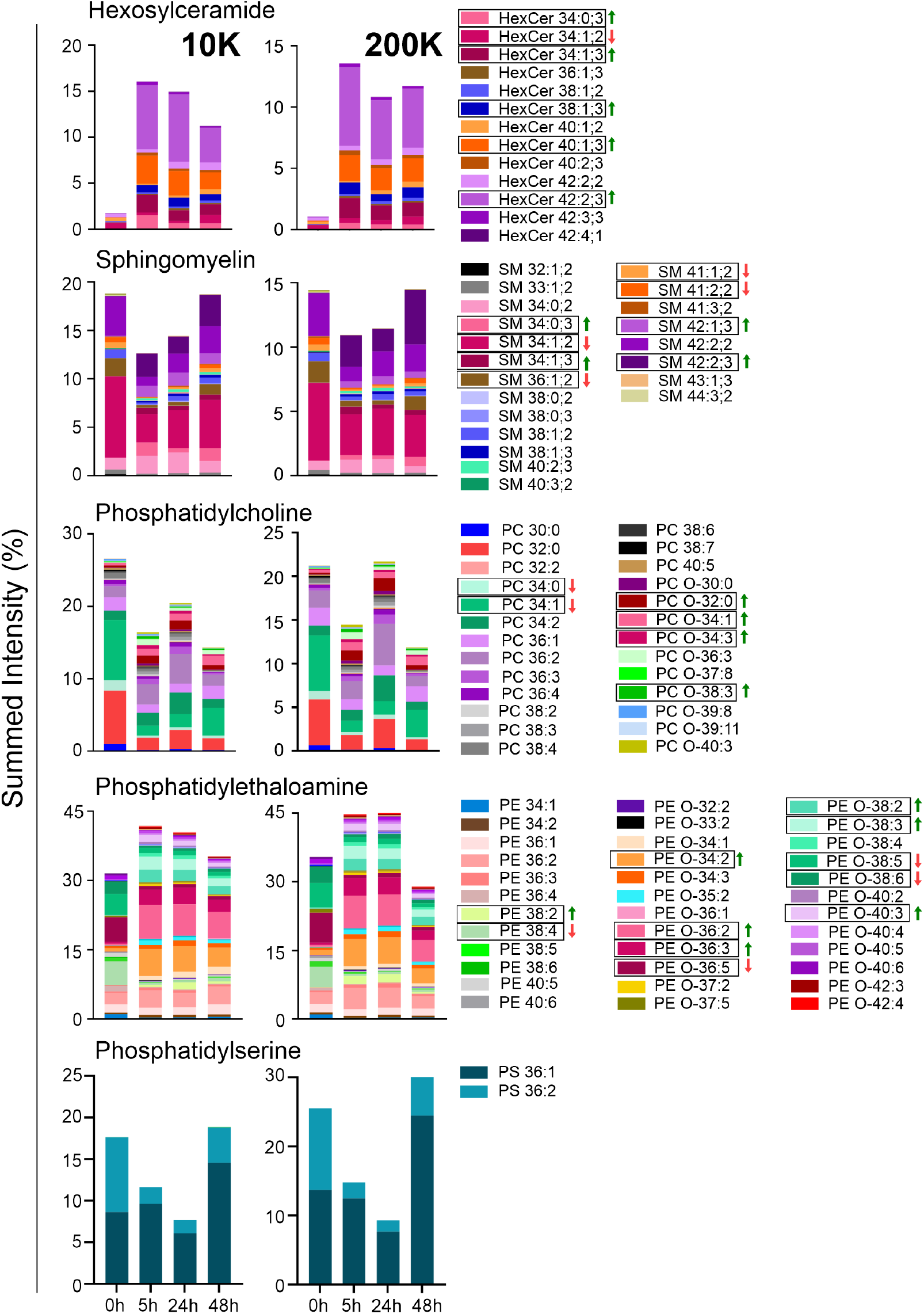
Composition of individual lipid species of SM, HexCer, PC, PE, and PS classes. Lipid composition of five lipid classes (HexCer, SM, PC, PE, PS, as labeled above the graphs) of SF–EVs at 0 hours (pre–LPS), 5 hours, 24 hours, and 48 hours (post–LPS). The black boxes highlight examples of lipids species that either increased or decreased in relative amount after the LPS stimulus compared to the baseline–healthy SF. A green arrow represents a relative increment of lipid species, and a red arrow a relative reduction. Lipids were obtained from 10k (left stacked bars) and 200k (right stacked bars) SF–EVs purification with sucrose density gradients. Healthy samples were taken pre–LPS–0h (n=4), synovitis samples at post–LPS–5h (n=4), synovitis–resolution samples at post–LPS–24h (n=3), and resolution samples at post–LPS–48h (n=3). Abbreviations: Hex2Cer (di–hexosylceramide); HexCer (hexosylceramide); PC, (phosphatidylcholine); PE, (phosphatidylethanolamine); PS, (phosphatidylserine); SM, (sphingomyelin).

### 2.3. Collection of equine synovial fluid

Synovial fluid was harvested by sterile arthrocentesis. Synovial fluid was centrifugated at 2,697xg (6,000 rpm; Centrifuge – Hettich EBA–20. Rotor 67 mm) for 30 minutes at RT to pellet cells and aggregates. The supernatant was stored at –80 °C for later EV isolation in aliquots of 1 ml (In vivo studies 1–4) or 7 ml (donated patients).

### 2.4. Extracellular vesicle isolation

A slightly modified protocol from our group was used to isolate EVs from SF [24]. In short, cell–free SF samples stored at –80°C were thawed (1 mL for each condition of the experimental horses and 7mL for the donated horses) and incubated in a water bath at 37°C for 15 minutes with 20 *μ*L HYase (5 mg/ml; Hyaluronidase (HYase) type II from sheep testes (≥300 units/mg), from Sigma–Aldrich (St. Louis, MO, USA)), 80 *μ*L DNase I (5 mg/ml; DNase I – grade II from bovine pancreas from Roche (Basel, Switzerland)), and 100 *μ*L sodium citrate (1.1 M; tri–sodium citrate dihydrate, from Sigma–Aldrich (St. Louis, MO, USA)) per 1 mL of SF. In order to remove (protein)aggregates, the SF was spun at 1,000xg for 10 minutes at RT (Eppendorf 5415R with rotor F45–24–11, Hamburg, Germany). Next, the supernatants were transferred into SW60 tubes (in vivo studies 1–4) or SW40 tubes (donated patients) (Beckman Coulter Inc., Brea, USA) and centrifuged at 10,000xg for 35 minutes, followed by a centrifugation at 200,000xg for 120 minutes. The SW 60 Ti and SW 40 Ti Beckman Coulter rotors were used in an Optima™ XPN–80 *μ*L tracentrifuge. EV pellets were resuspended in 300 *μ*L PBS/0.1% BSA (depleted from EVs and protein aggregates).

EV pellets were mixed with 1.2 mL sucrose solution 2.5 M in PBS (J.T. Baker, Phillipsburg, NJ, USA) at the bottom of SW40 ultracentrifuge tubes. Subsequently, fourteen sucrose solutions of decreasing molarity were carefully overlaid (from 1.88 M to 0.4 M) to create discontinuous sucrose gradients. Gradients were centrifuged at 200,000xg for 16 hours at 4°C (Optima™ XPN–80 centrifuge; SW40 Ti rotor; 40,000 rpm; relative centrifugation force (RCF) average 202,048xg; RCF max 284,570xg; *κ*–factor 137; Beckman–Coulter, Fullerton, USA). After centrifugation, 12 fractions of 1 mL were collected from the top (fraction 12) to the bottom (fraction 1). Fraction densities were assessed by refractometry. See Suppl. Fig. 1 for a schematic representation of the EV–isolation procedure.

### 2.5. Extracellular vesicle characterization

#### 2.5.1. Single–EV high–resolution flow cytometry (FCM)

Labeling of EVp with PKH67. Generic staining was performed as previously described [25] with minor modifications as described below. EV pellets were resuspended in a mixture of 20 *μ*L EV–depleted PBS/0.1% EV–depleted BSA and 30 *μ*L diluent C. Next, 50 *μ*L diluent C was added and mixed with 1.5 *μ*L of PKH67 dye. After 3 min incubation at RT, the staining reaction was stopped by adding 50 *μ*L of EV–depleted RPMI/10% FBS. Labeled EVs (250 *μ*L) were then mixed with 2.5 M sucrose for the continuation of EV isolation with density gradient ultracentrifugation as described in section 2.4.

A control sample (20 *μ*L EV–depleted PBS/0.1% EV–depleted BSA+30 *μ*L diluent C, without EVs) was taken along during the entire PKH67 labeling and sucrose gradient ultracentrifugation procedure. The high–resolution FCM of this control sample did not show significant background events in gradient fractions of interest (data not shown). *Single*–*EV*–*based high*–*resolution FCM*. An optimized jet–in–air–based flow cytometer (BD Influx, Beckton Dickinson Biosciences, San Jose, CA, USA) was used, as previously described [25; 26]. Briefly, a workspace with pre–defined gates and optimal PMT settings for the detection of 200 nm yellow–green (505/515) FluoSphere beads (Invitrogen, F8848) was loaded. Upon aligning the fluid stream and lasers, the 200 nm bead population had to meet the criteria of pre–defined MFI and scatter values within these gates, displaying the smallest coefficient of variation (CV) for side scatter (SSC), forward scatter (FSC) and FL–1 fluorescence. The trigger threshold level was kept identical for all measurements in this study and set by running a clean PBS sample, thereby allowing an event rate of ≤10–20 events/second. All samples were acquired for a fixed time of 30 seconds. EV concentration was calculated by measuring the instrument’s flow rate and correcting the number of detected particles for dilution and measured time. The EV concentration per ml of SF was calculated by taking the sum of EVs in sucrose fractions F3–F9, corrected for the SF starting volume.

Data were acquired using BD FACS™ Software v1.0.1.654 (BD Biosciences, San Jose, CA, USA) and analyzed with FlowJo v10.07 software (FlowJo, Ashland, OR, USA).

#### 2.5.2. Cryo–TEM

Preparation of samples. EV pellets were resuspended in 250 *μ*L PBS, mixed with 1.25 ml 2 M sucrose solution (in PBS) (total volume = 1.5 ml) in MLS50 tubes, and carefully overlaid with sucrose solutions of 1.4, 0.4, and 0 M (sucrose in PBS) to create 2 small discontinuous sucrose gradients. Gradients were centrifuged at 200,000xg for 16 h at 4°C (Beckman–Coulter Optima™ MAX–E centrifuge; MLS50 rotor; 50,000 rpm; RCF average 200,620xg; RCF max 268,240xg; *κ*–factor 71.1). After centrifugation, 5 fractions of 1 ml were collected from the bottom (F5) to the top (F1) using a peristaltic pump. Densities were determined by refractometry. EV–containing fractions were pooled (1.10–1.18 g/mL) and diluted by adding 9 ml PBS to SW40 tubes (Beckman–Coulter, Fullerton, CA, USA). EVs were pelleted at 200,000xg for 60 min at 4°C (Beckman–Coulter Optima™ L–90K or Optima™ XPN–80 centrifuge; SW40–Ti rotor; 39,000 rpm; RCF average 192,072xg; RCF max 270,519xg; *κ*–factor 144.5). *Cryo*–*TEM*. EV pellets were carefully resuspended in 20 *μ*L PBS and stored on ice for 1–2 h until vitrification using a Vitrobot™ Mark IV system (FEI, Eindhoven, Netherlands). Next, three microliters of EV sample were directly placed onto a glow–discharged 2/2 copper grid (Quantifoil, Jena, Germany). The excess sample was removed with 595 filter paper (Schleicher & Schuell, Whatman Plc., Kent, UK) in the Vitrobot chamber for 1 s at 100% relative humidity, with subsequent plunging into liquid ethane (3.5 purity). Residual ethane was removed with filter paper, and grids were stored in cryo–boxes under liquid N2 for later imaging. For cryo–TEM, grids were transferred to a Gatan 626 cryo–holder (Gatan Inc., Pleasanton, CA, USA) and inserted into a Tecnai™ 20 transmission electron microscope (FEI) with LaB6 filament operated at 200 kV. Images were acquired with a 4,000×4,000 Eagle charge–coupled device (CCD) camera (FEI) at a 19,000× magnification, 5–10 um under focus.

#### 2.5.3. Western blotting

For Fig. 1D (Suppl. Fig 2), sucrose fractions were pooled in pairs (1.06–1.08 g/mL, 1.10–1.12 g/mL, 1.14 – 1.16 g/mL) and diluted in PBS in SW40 tubes. For Suppl. Fig. 3B, individual fractions were diluted in PBS in SW60 tubes. Sucrose fractions were spun at 200,000xg for 90 min at 4°C. EV pellets were suspended in 65 *μ*L non–reducing SDS–PAGE sample buffer (50mM TRIS pH 6.8, 2% w/v SDS, 10% v/v glycerol, 0.02% w/v bromophenol blue), heated at 100°C, run on 4–20% Criterion TGX gels (Bio–Rad, Hercules, CA, USA), and transferred onto 0.2 um PVDF membranes (Bio–Rad, Hercules, CA, USA) or 0.22 um nitrocellulose membranes (MilliposeSigma, Burlington, MA, USA). After blocking for 1 hour in blocking buffer (for Fig. 1D: 5% Protifar; for Suppl. Fig. 2B: 0.2% v/v Fish skin gelatin (Sigma–Aldrich, St. Louis, MO, USA)+ 0.1% v/v Tween–20) in PBS, proteins were labeled with primary antibodies against CD9 (1:1,000; clone HI9, Biolegend, San Diego, USA), CD44 (1:400; clone IM7, PE–conj., eBioscience, San Diego, USA), MHC Class II (1:400; clone CVS20, Bio–Rad, Hercules, CA, USA) and Annexin–A1 (1:400; clone 29/Annexin–1, BD Transduction Labs, San Jose, CA, USA). HRP–conjugated goat–anti–mouse IgG (1:10,000; Nordic Immunology Laboratories, Susteren, the Netherlands) was used for detection by chemiluminescence. Chemiluminescence was visualized using a ChemiDoc™ MP Imaging System (Bio–Rad, Hercules, CA, USA) and analyzed with Bio–Rad Image Lab V5.1 software (Bio–Rad, Hercules, CA, USA).

### 2.6. Phospholipid analysis

#### 2.6.1. Lipid extraction

Lipids were extracted according to the Bligh & Dyer method [27] with slight modifications. First, 0.8 mL was taken from each sucrose fraction and mixed with 1 mL chloroform and 2 mL methanol. After 20 minutes, 2 mL chloroform and 2 mL of H2O were added, and after thoroughly mixing, the two phases were separated by centrifugation for 5 minutes at 2,000xg. Next, the bottom phase – the chloroform phase containing the lipids – was transferred to another conical glass tube, and 2 mL of chloroform was added to the remaining sample (upper phase), which was again vortexed and centrifuged. Afterward, the bottom chloroform phase was once more collected and pooled with the first chloroform extract. Finally, the samples were dried under nitrogen gas injection, and the dried pellets were stored in a nitrogen atmosphere at –20 °C.

#### 2.6.2. Ultra–high performance liquid chromatography – Mass Spectrometry (UHPLC–MS) of lipids

The dried lipid extract was dissolved in 20–30 *μ*L chloroform/methanol (1:1) and analyzed as reported previously [28]. Ten microliters of lipid extract were injected into a hydrophilic interaction liquid chromatography (HILIC) column (2.6um Kinetex HILIC 100Å, 50×4.6mm, Phenomenex, Torrance, USA). Lipid classes were separated by gradient elution using an Agilent 1290 InfinityII UPLC (Agilent, CA). Solvent A consisted of acetonitrile/acetone (9:1) with 0.1% formic acid, and solvent B consisted of acetonitrile/H2O (7:3) with 0.1% formic acid and 10 mM ammonium formate. The gradient elution was: minute 0 to 1 – 100% A; minute 1 to 3 – 50% A + 50% B; minute 3 to 5 – 100% B. The flow rate was 1 mL/minute. Successive samples were infused without further re–equilibration of the column.

Afterward, the samples entered into a Fusion Orbitrap MS (ThermoFisher Scientific, Waltham, USA) – in a negative ion mode – where they underwent a heated electrospray ionization (HESI) with the following conditions: vaporizer temperature, 450°C; ion transfer tube temperature, 350°C; negative ion spray voltage, 3.6 kV; sheath gas flow rate, 54Arb; aux gas flow rate, 7Arb; sweep gas flow rate, 1Arb; scan range, 350–950 m/z at a resolution of 120,000.

### 2.7. Data analysis

The data were converted to mzML format by msconvert ProteoWizard [29] using the vendor peak picking filter. LC/MS peak–picking, sample–grouping, and rt–correction of the mzML–files were performed using XCMS version 3.10.2 [30–32] running under R version 4.0.3 [33]. Using an in silico phospholipid database, the recognized LC/MS peaks (features) were annotated based on retention time, exact m/z–ratio, and, if present, MS2 spectra of the features. Moreover, the features were corrected for isotope distribution and adducts. The data were normalized by expressing all lipid species as fractions of the sum of the total annotated lipid intensity of each biological sample. Features were furthermore selected according to the consistency within two technical replicates. Statistical analyses were based on at least three biological replicates; if the lipid levels of a data point were below detection, these were omitted from further analysis.

Fig. 1B, C, E, and 5, and Suppl. Fig. 3A, C, and 4 were generated with Prism version 9.1.2 (© 1992–2021 GraphPad Sofware, LLC). Partial least squares discriminant analysis (PLS–DA) was performed on normalized data using the mixOmics R package version 6.12.2. [34]. Heatmap and cluster analysis was performed on Spearman correlations with a column k–means algorithm of three and a set.seed of two – among the 100 most abundant lipid species in all sample groups – using the R–package ComplexHeatmap v1.12.0 [35].

## 3. Results

### 3.1. Equine synovial fluid–derived EV analysis after LPS injection

To analyze whether the number and composition of SF–derived EVs changes during joint inflammation, synovitis was induced by LPS injection in equine joints, and EVs were isolated from SF before and after LPS injection. Previously, we reported and validated a protocol for the isolation of EVs from equine SF derived from healthy joints, which we applied in the current study [24]. We first performed a morphological analysis by cryo–EM of SF–EVs isolated during the acute inflammation phase (at 8h) after pelleting by 10K or 200K, followed by sucrose density gradient centrifugation. For the 10K EVs, 197 vesicles with heterogeneous shapes and sizes were analyzed, while for the 200K, 485 were analyzed (Fig. 1A, 1B). There were no significant variations in the percentage of spherical EVs for the 10K and 200K of the 8h–synovitis SF–EVs vs. the data from healthy joints as reported previously [24] (82% of 10K and 53% of 200K 8h–synovitis SF–EVs, against 65% and 67% in the healthy sample, respectively). This is in contrast to the amount of “other small particles” which drastically increased in the 8h–synovitis 200K SF–EVs to 25% compared to 3% in the healthy sample. Regarding the size, healthy SF–EVs had a slightly bigger average size of spherical vesicles compared to synovitis SF–EVs, with an average size of 109 nm for 10K and 90 nm for 200K of healthy SF–EVs vs. 90 nm for 10K and 78 nm for 200K of 8h–synovitis SF–EVs (Fig. 1C). Importantly, the size distribution of 10K and 200K EVs did not substantially differ (Fig. 1C).

Western blot analysis on pooled EV–enriched densities validated the expression of the tetraspanin CD9 (an EV marker) in healthy SF–EVs and 8h–synovitis SF–EVs. Equally important, three other EV–associated proteins, CD44 (HYase–receptor), AnxA1 (anti–inflammatory protein), and MHC–II (a marker for antigen–presenting cells) were all readily detectable in the 8h SF–EVs samples (Fig. 1D, Suppl. Fig 2).

To determine the effect of LPS–induced synovitis on the amount of SF–derived EVs, we performed single vesicle fluorescence–based flow cytometric analysis of EVs using a previously optimized method [25; 26]. Hereto, SF–EVs isolated before LPS injection (0h) or after LPS injection during acute inflammation (8h) and during the resolution phase (24h and 168h) were labeled with PKH67 and analyzed (Fig. 1E). At 8h during acute inflammation, there is a sharp increase of fluorescent events compared to 0h SF–EVs both for 10K and 200K EVs. However, at 24h post–LPS injection, the numbers had decreased for both 10K and 200K EVs, which persisted after 168 hours when fluorescent events had returned to baseline. Subsequently, the same kinetics were observed in another horse study in which SF was collected at 0h, 5h, 24h, and 48h after interarticular LPS injection and corroborated by the CD9 western blot data of the corresponding samples (Suppl. Fig. 3A–B). Hence, we demonstrate that during LPS–induced synovitis, the numbers of EVs rapidly increase in the acute phase (5–8h) and start to decline at 24h post–LPS injection and return to healthy joint baseline levels between 48–168h after LPS injection.

### 3.2. The effect of acute LPS–induced synovitis on the phospholipidome of EVs isolated from equine synovial fluid

#### 3.2.1. Relative total phospholipid intensity

To investigate the effects of LPS as an inducer of inflammation on the SF–EVs lipidome, SF samples of three horses that underwent an LPS injection and three horses with healthy joints were employed to decipher the phospholipidome of SF–EVs. Results showed that there was an increase of lipids in the samples collected after the experimentally induced inflammation (Suppl. Fig. 3C), which correlates with the EV dynamics discussed in section 3.1. Since the numbers of EVs (Fig.1E and Suppl. Fig.3A), and thus lipids (Suppl. Fig.3C), are much lower in healthy SF vs. Synovitis SF, equal input volumes of SF resulted in many lipids below detection levels in 2 out of 3 healthy SF–derived EV samples (Suppl. Fig.3C). Therefore, 3 additional healthy SF–derived EV samples were prepared from SF collected from the donated horses with healthy joints from which seven times more SF could be harvested. The samples shown in Supplementary Fig. 3C and the 3 additional healthy SF–EV samples were normalized for each time–point (0h, 5h, 24h, and 48h) and EV–containing sucrose fraction (F6 to F10 corresponding to densities from 1.18 g/mL to 1.10 g/mL) by expressing the lipid intensity as a fraction of the sum of the total phospholipid intensity, hence identifying the peak densities in all samples irrespective of their absolute lipid levels (Suppl. Fig. 4). This approach showed that the peak densities were different depending on the original EV pellet, with the peak of 10K SF–EVs in densities 1.16–1.14 g/mL and for 200K SF–EVs in densities 1.14–1.12 g/mL. For the following detailed phospholipid analysis, the SF–EVs of 10K and 200K EVs, present in density fractions 1.16–1.14 g/mL and 1.14–1,12 g/ml, respectively, were analyzed independently.

#### 3.2.2. Alteration of SF–EVs lipid profile

Elucidation of the composition of the SF–EV phospholipidome was achieved by a bioinformatics analysis which revealed 158 lipid species after lipid annotation (and background adjustment), isotope and adduct correction, normalization, peaking of consistent features between technical replicates, and selection of the peak densities for 10K and 200K separately (Suppl. Fig. 4–6). Subsequently, the lipidomic data was visualized with partial least square discriminant analysis (PLS–DA) for 10K and 200K SF–EVs (Fig. 2). The scatter plot grouped the samples according to the two first discriminants. The PLS–DA illustrated that healthy SF–EVs had a significantly different lipid profile in comparison to the post–LPS injection SF–EVs. Furthermore, it was noticed that the 5h and 24h SF–EVs samples largely overlap. It is also clear that the group of 48h EVs is closer to the 0h EVs. From this, we can infer that the lipidome slowly returns to baseline 48 hours after the LPS stimulus. It should be noted that a shift of the same group on the PLS–DA2 axis indicates a change in the lipidome of the 48h EV group relative to the other groups. In general, the patterns for the profiles of 10K and 200K EVs are rather similar, suggesting that the phospholipid profiles of these different EV samples are similarly affected after LPS injection.

To better illustrate the distribution of the lipids for the two EV populations (10K and 200K), a correlation heatmap of the 100 most abundant lipid species was constructed (Fig. 3). Both heatmaps show two distinct clusters (Fig. 3A and 3B). Going from left–to–right, in the first slice of both the 10K EV (Fig. 3A) and 200K EV heatmap (Fig. 3B), the two display clusters consist of lipids prevalently present (according to the group distribution) in the post–LPS–5h SF EVs (red) and post–LPS–24h SF EVs (orange) and to some extent in the post–LPS–48h SF EVs (blue). These lipids are prominently present in the inflammatory profile, as indicated in our proposed interpretation. In the center of the heatmap (between the white arrows), there are no clearly delineated clusters, pointing towards a more random group distribution. On the right side of the heatmap, two defined clusters are displayed, depicting lipids prevalent in healthy SF EV samples (green, 0h samples). The results from Fig. 3A and 3B are in line with the PLS–DA loading plots, in which the 5h and 24h SF–EVs were closely related, and the 0h SF–EVs had a different profile. The 48h SF–EVs showed a slight similarity with the 0h samples but were still closely associated with the 5h and 24h SF–EVs.

#### 3.2.3. Variations in the lipid class composition of SF–EVs

To analyze the effect of LPS injection on the lipidome profile of SF–EVs in greater detail, we next evaluated the fluctuation in lipid class compositions. Figure 4 represents a fractional composition plot depicting the fluctuation of the lipid class compositions of SF–EVs among the four different time points. The induction of inflammation (post–LPS–5h) leads to a relative increase in ether phosphatidylethanolamines (PE O–, including plasminogens) and a sharp upsurge of hexosylceramides (HexCer, i.e., glucosyl– or galactosylceramides) for both the 10K and 200K SF–EVs. At the same time, phosphatidylserines (PS) and sphingomyelins (SM) relatively declined in EVs from both 5h and 24h post–LPS compared to 0h (healthy) and 48h post–LPS EVs. Ester–linked PE likewise slightly decreased after inflammation induction, however, these lipids were not back to healthy (0h) levels in 48h post–LPS EV samples. Finally, phosphatidylcholine showed a complex dynamic fluctuation, in which the 0h and 24h post–LPS SF–EVs have relatively more PC than the 5h and 48h SF–EVs. Taken together, we here identified inflammation–related fluctuations in the lipid classes of EVs, with healthy SF–EVs and 48h SF–EVs having relatively similar profiles, while synovitis SF–EVs (5h and 24h post–LPS) present a different makeup.

#### 3.2.4. Turnover of key SF–EVs lipid species after inflammation

We next determined which individual species of each of the five predominant lipid classes contributes to the fluctuations in the EV–lipid classes during the course of inflammation (Fig. 5). In Fig. 5, single lipid species – key in the lipid profile fluctuation – present within SF–EV samples isolated at different time points are indicated. The hexosyl-ceramide (HexCer) class for both 10K and 200K expands greatly after the LPS insult. Most lipids species increase in intensity, for instance, HexCer 42:2;3, HexCer 38:1;3, HexCex 40:1;3, and HexCer 34:1;3. In contrast, HexCer 34:1;2 shrinks in the synovitis SF–EVs to increase again in the 48h SF–EVs. Inside the sphingomyelin (SM) class, some lipids rise in relative proportion after the inflammatory stimulus – e.g., SM 42:2;3, SM 34:0;3, SM 34:1;3, and SM 42:1;3 – and other lipids decreased in proportion, such as SM 42:2;3, SM 34:0;3, SM 34:1;3, and SM 42:1;3. The phosphatidylcholine (PC) class lipids exhibit a peculiar but consistent pattern in both the 10K and 200K SF–EVs. As with the SM class, the PC general class profile is made of similar lipid species, with given species increasing after the inflammatory stimulus – e.g., PC–O 32:0, PC–O 34:1, PC–O 34:3, PC–O 38:3 – and others declining – e.g., PC 34:0, PC 34:1. Conversely, the most salient observation is how the total amount of phosphatidylcholines seems reduced at 5h, goes up at 24h, and declines again at 48h. This effect is consistent in both 10k and 200k EV preparations and is caused by certain specific ester PC lipids – notably PC 30:0, PC 32:0, PC 34:0, PC 34:2, PC 36:2, PC 36:3, and PC 36:4. In the phosphatidylethanolamine (PE) class (composed of ester– and ether–linked PE) there is more disparity. This class has the highest diversity of lipids species. In this class, too, some species rise after the LPS injection (like PE 38:2, PE–O 34:2, PE–O 36:2, PE–O 36:3, PE–O 38:2, PE–O 38:3, and PE–O 40:3) and others relatively decrease (PE 38:4, PE–O 36:5, PE–O 38:5, and PE–O 38:6). The color profile of the four PE stacked bars illustrates how different the PE profile is in all post–stimulus groups compared to baseline. Lastly, the phosphatidylserine (PS) class is made up of only two lipids. In all conditions, PS 36:1 was the dominant PS. Nevertheless, the ratio of both PS, which is roughly equal in the healthy state, changes substantially due to the deep drop in PS 36:2 after the inflammatory stimulus that decreases the entire class. At 48h, the overall drop has been compensated for, but largely through an increase in PS 36:1, leaving the ratio skewed in comparison to healthy SF EVs.

## 4. Discussion

Thus far, EV research in joint diseases has primarily focused on the use of EVs as a diagnostic tool or a potential therapeutic for several types of pathologies. Most of those studies concentrate on the protein and nucleic content of EVs, largely disregarding the potential role of the surrounding lipid membrane. Comprehending the dynamics of the large lipid fraction of EVs in health and disease may, then again, be essential to fully understand the functionality of these structures. To partially fill this gap, the current study aimed at characterizing, quantifying, and analyzing EVs from synovial fluid (SF–EVs) after the induction of an acute and marked, yet transient, inflammation of the equine joint through the interarticular injection of bacterial lipopolysaccharide (LPS).

The morphology of the SF–EVs from healthy joints [24] and synovitis joints was observed to be vastly similar. The few morphological discrepancies noted were a slightly smaller size of SF–EVs and an increase in the number of small particles other than EVs in inflamed joints. On the contrary, EV dynamics were strikingly different, with SF from a healthy state showing much fewer EVs and lower lipid intensities in comparison to the SF EVs isolated at the peak of inflammation (5h/8h). This steep upsurge after the LPS insult was followed by a noticeable reduction of EV levels already at 24h, which further declined until healthy SF baseline levels at 168h. This outcome shows that EV release from resident and/or infiltrated cells peaks at five to eight hours after induction of acute synovitis. These results are in agreement with previous findings on EVs in joint inflammation [10; 36] and with the relatively high EV concentrations found in patients with arthritis [37–39]. The results also indicate that, if measured with single EV–based analysis techniques, e.g., the high–resolution flow cytometry used in the current paper, or even with straightforward mass spectrometry analysis of phospholipids, the EV concentration in SF could potentially already be used as a biomarker for the presence of joint inflammation. The impact of an inflammatory insult on the SF–EVs in the joint was further assessed by western blot analysis of EV–associated markers that could be detected in equine samples. At the peak of inflammation (8h after the LPS insult), besides the EV marker CD9, we detected CD44, AnxA1, and MHC class II on SF–EVs. Both CD44 and AnxA1, albeit distributed on several different cell types, are known to be expressed on neutrophil–derived EVs [38; 40–42], the predominant inflammatory cell type in acute LPS–induced arthritis [21; 38; 41; 42]. Interestingly, in these studies, neutrophil–derived EVs have been associated with an anti–inflammatory role [38; 41]. On the other hand, we also demonstrated the presence of MHC–II+ EVs at 8h in the 200K EVs. These EVs probably arise from antigen–presenting cells and may be involved in driving T–cell responses [43; 44]. Overall, the role of SF–EVs during inflammatory joint disease processes is still poorly understood.

We here showed that the SF–EV lipidome profile reacted heavily to the inflammatory stimulus. Overall, SF–EVs isolated 5h and 24h post–LPS injection have rather similar lipid profiles that are considerably different from healthy SF–EVs. At 48h, the lipidic architecture subtly begins to resolve back to a healthy state. This pattern is congruent with previously observed patterns of inflammatory markers in LPS–induced synovitis studies, such as prostaglandin E2 (PGE2), bradykinin, substance P, MMP activity, interleukin–1, and tumor necrosis factor–*α* (TNF*α*) [21; 45]. These differences in lipidomic profile arise from the hydrophilic head, hence the lipid class, of which we focused on the top five most abundant: HexCer, SM, PS, PE, PE O– and PC. Ceramides (Cer) were disregarded in the analysis due to their retention time near the solvent front. Nevertheless, even if we could not determine the changes in ceramides directly, we were able to observe how one of its metabolites (HexCer) significantly increased during acute inflammation. Interestingly, higher levels of circulating HexCer have been associated with inflammatory diseases (e.g., rheumatoid arthritis and multiple sclerosis) [46–48], and high levels of HexCer can play a crucial step in the production of TNF*α* as part of an immune response cascade that leads to the activation of COX2 [49] a precursor activator of PGE2 – a well–known inflammatory lipid mediator [17]. Besides, HexCer can serve as a substrate for the synthesis of more complex glycosphingolipids [50]. Therefore, we hypothesize that the SF–EVs present at 5h, 24h, and 48h post–LPS injection, which are substantially enriched in HexCer, are carriers of HexCer and possibly also Cer to cope with the high demand for lipids necessary for the upscaling of the biochemical cascade leading to the formation of lipid–based inflammatory mediators such as prostaglandins. In addition, it has been established that TNF*α* activates nSMase–2. The role of nSMase–2 is the hydrolyzation of SM to transform it into Cer, hence aiding its accumulation [51]. At the same time, this leads to the reduction of available SM. We indeed saw a decrease in SM levels at 5h and 24h during the inflammation peak. However, the levels of SM were restored at 48h at the onset of the resolution phase. These results on HexCer and SM are congruent with the presumed role of sphingolipids as structural components of plasma membranes and bioactive molecules with an essential role in proliferation, differentiation, growth, signal transduction, and apoptosis [50; 52; 53].

Moreover, HexCer and SM have been reported to be components of lipid rafts, which aid in the selection of membrane proteins associated with signal transduction and intracellular transport [54; 55]. Lipid rafts have also been linked to exosome formation [56–58]. Exosome biogenesis requires Cer, which is produced after SM hydrolysis by nSMase [59], and HexCer by ceramide glucosyltransferase [60]. In fact, vesicular trafficking encourages the conversion of Cer into HexCer [61]. Therefore, it is not unlikely that HexCers also function in the assembly of EVs. In fact, it has been proven that Lysosome Associated Protein Transmembrane 4B (LAPTM4B) plays a role in the homeostasis of HexCer by suppressing its presence – and of its downstream metabolite, globotriaosylceramide (Gb3) – in both cells and EVs [62]. Also, SM and Cer collaborate in exosome biogenesis through the ESCRT–independent pathway, forming intraluminal vesicles inside multivesicular bodies (MVBs), which are later released as exosomes after MVB fusion with the plasma membrane [59; 63].

As for the ester–linked PE, we noticed a decrease in SF–EVs isolated after the LPS stimulus. The levels continue to be low, even at 48h. That aside, a more remarkable effect was observed within the ether–linked PE where a rise after the LPS injection occurred, which level remained stable at 24h, but dropped at 48h to come closer to baseline levels seen in healthy SF–EVs. Ether–linked lipids (including plasminogens) have single–bonded oxygen (annotated as PX O–). Overall, we detected lower amounts of ester PE versus ether PE in all conditions. Thus, ether PE is preferred by the cell while building the EVs, probably for their predisposition to take on non–lamellar inverted hexagonal shapes, enabling membrane fusion processes. Also, ether lipids are involved in lipid raft microdomains and take part in membrane trafficking, cell signaling, and differentiation [64; 65]. Furthermore, ether–linked lipids allow a compact packing of phospholipids in the membranes, thus lowering membrane fluidity and making these more rigid [66]. Enriched ether–linked membranes have also been found in several cells and tissues, such as the brain, heart, kidney, lung, skeletal muscle, eosinophils, and neutrophils [67; 68].

At 5h and 24h post–LPS injection, when we observed an increase in ether–linked PE in SF–EVs, the amount of PS was reduced. It is worth noting that PE and PS are lipid classes found primarily in the inner leaflet of the lipid bilayer of cell plasma membranes and EVs [69; 70]. There are data suggesting that PS 36:1 has a close interaction with outer long–chain lipids, such as SM, producing a type of hand–shaking interaction [71], which would result in greater cohesiveness and rigidity of the EV membrane. This could explain why PS 36:2 was preferentially depleted, given the critical role of PS 36:1, which likely is co–sorted with the fellow long–chain sphingolipids [71]. Moreover, PS 36:1 has been shown to prevent oxidation by withholding cholesterol – a crucial participant in maintaining the membrane fluidity and incrementing the lipid–density packaging – in the inner leaflet of the membrane [72; 73]. PS 36:1 is as well known to interact with and bind to cytoplasmic proteins (such as members of the Rab family), activating a number of signaling cascades [74]. PS has also been associated with processes concerning the dynamics of the plasma membrane, such as the assembly of caveolae [75]. At 48h, PS overall returns to the same level as at 0h. Nevertheless, the profile of both PS species shows a substitution between the two PS lipids. With a 1:1 ratio in a healthy state, which is shifted during inflammation with a sharp drop in PS 36:2, it is, at 48h when the overall level of PS has been restored, still PS 36:1 that compensates for the lack of PS 36:2, proving the importance of PS 36:1.

In contrast to the other lipid classes, the SF EV–associated PC class depicts a less consistent pattern with varying levels throughout the different phases after LPS injection. The effect is caused solely by seven specific ester–linked PC species (PC 30:0, PC 32:0, PC 34:0, PC 34:2, PC 36:2, PC 36:3, PC 36:4). Unfortunately, it is uncertain what the specific roles of those PC species are.

All in all, our results demonstrate a profound effect of acute inflammation on synovial fluid–derived extracellular vesicles (Suppl. Fig. 6). We found an upsurge in EV dynamics and substantial transformation of the lipidome profile regardless of the vesicle size (i.e., 10K EVs vs. 200K EVs), which takes longer than 48h to restore itself to the healthy profile at the lipid level. The observed changes in the lipidome are likely due to the presence of new EVs from different cell populations that migrate toward the joint cavity in response to the inflammatory stimulus or from an altered EV profile released by synovial lining cells. Therefore, further work is needed on characterizing the EV lipidome of specific cell types that infiltrate the joint and the modulation of synovial lining cells during various forms of synovitis, such as during RA or OA. Importantly, in this study, we identified specific lipidomic profiles associated with healthy or inflammatory state–derived SF–EVs and pinpointed key lipid species that were critical for the lipidome turnover at different time points.

## Acknoledgments

We thank Willie J. C. Geerts and Johannes Bergmann (former master student) for performing cryo–TEM; Filipe M. Serra Bragança, Janny C. de Grauw, and Stefan M. Cokelaere for supplying the synovial fluid material from their respective animal experiments for this study; Ger J. A. Arkesteijn for providing help with the single–EV high–resolution flow cytometry (FCM).

## Declaration of author contributions

**Marca H.M. Wauben:** Conceptualization, Methodology, Project administration, Supervision, Validation, Writing – Review & Editing; **P. René van Weeren:** Conceptualization, Funding acquisition, Project administration, Supervision, Writing – Review & Editing; **Chris H.A. van de Lest:** Conceptualization, Data Curation, Formal analysis, Validation, Visualization, Supervision, Software, Writing – Review & Editing; **Laura Varela:** Formal analysis, Investigation, Validation, Visualization, Writing – Original Draft, Writing – Review & Editing; **Janneke Boere:** Formal analysis, Investigation, Visualization, Writing – Original Draft, Writing – Review & Editing; **Sten F.W.M. Libregts:** Investigation, Writing – Review & Editing; **Estefania Lozano**–**Andrés:** Investigation, Writing – Review & Editing.

## Supplementary figures

**Figure 6: Suppl. Fig. 1.**
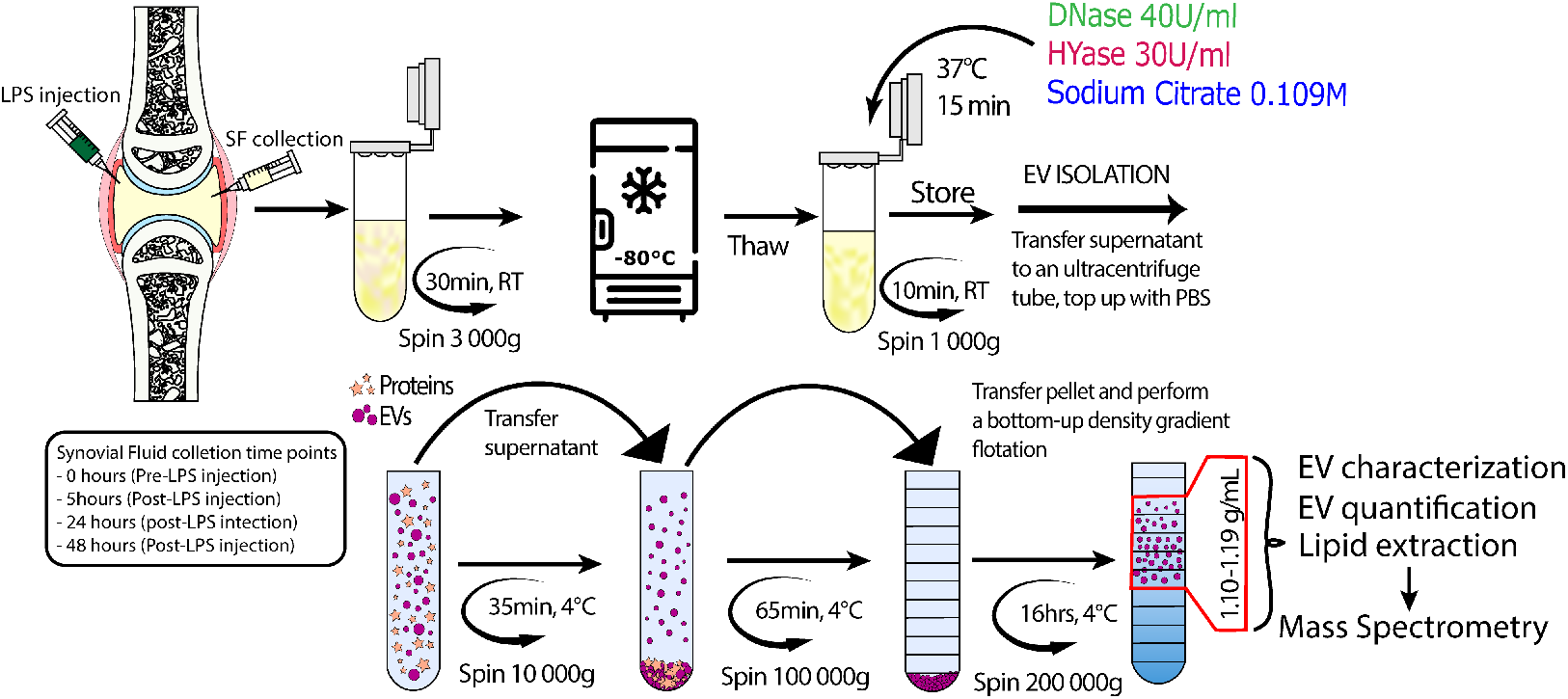
Schematic representation of the isolation protocol for extracellular vesicles from synovial fluid. Synovial fluid (SF) was collected pre-LPS injection from the middle carpal joints of Warmblood donors at 0 hours, followed by synovitis induction by intra-articular injection of lipopolysaccharide (LPS) from E. coli. Subsequently, SF was harvested at 5 hours, 24 hours, and 48 hours. SF samples were spun at 3,000xg for 30 minutes and stored at -80°C. Synovial fluid was thawed and treated with DNase (10U/ml), HYase (30U/ml), and EDTA (4.83mM) for 15 minutes at 37°C and spun at 1,000xg. EVs were isolated by differential centrifugation at 10,000xg (10K) and 200,000xg (200K). Extracellular vesicles were purified by sucrose density gradient floatation. Fractions 6 to 10 (1.10 – 1.18 mg/ml) were chosen for lipid extraction and subsequent lipidomic analysis. Densities of individual fractions (g/mL) are: F1=1.26, F2=1.25, F3=1.24, F4=1.22, F5=1.20, F6=1.18, F7=1.16, F8=1.14, F9=1.12, F10=1.10, F11=1.08, F12=1.06. Lipids were extracted by the Bligh & Dyer method and analyzed in the Orbitrap-Fusion MS in negative mode.

**Figure 7: Suppl. Fig. 2.**
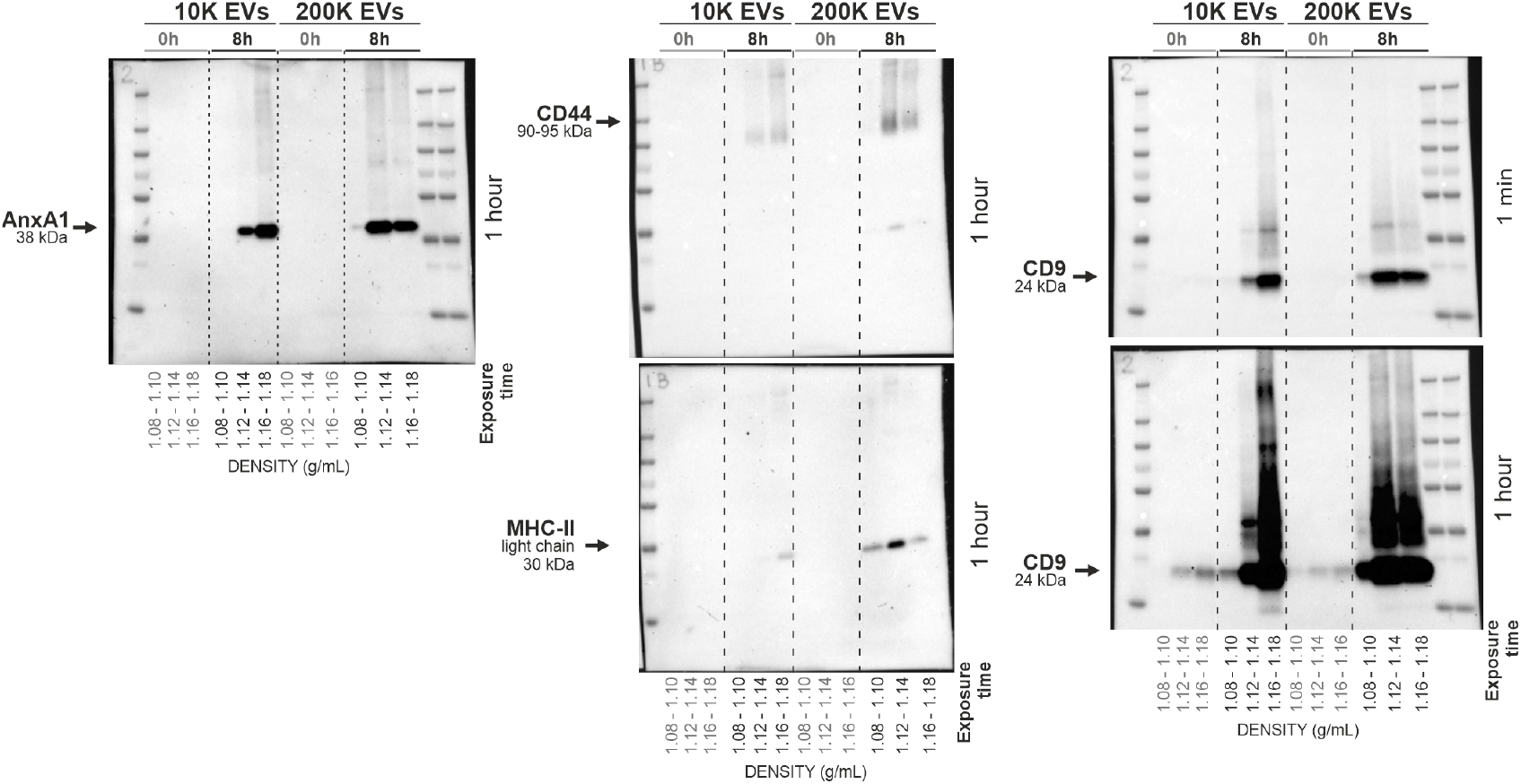
Full western blots from EVs isolated from joints with acute synovitis or healthy joints. Complete western blot analysis of EV-containing sucrose gradient fractions for the identification of CD9, CD44, AnxA1, and MHC-II proteins. Lanes were loaded with a pool of 2 sucrose fractions for 10k and 200k-derived EVs. The selected protein markers used various exposure periods of up to an hour.

**Figure 8: Suppl. Fig. 3.**
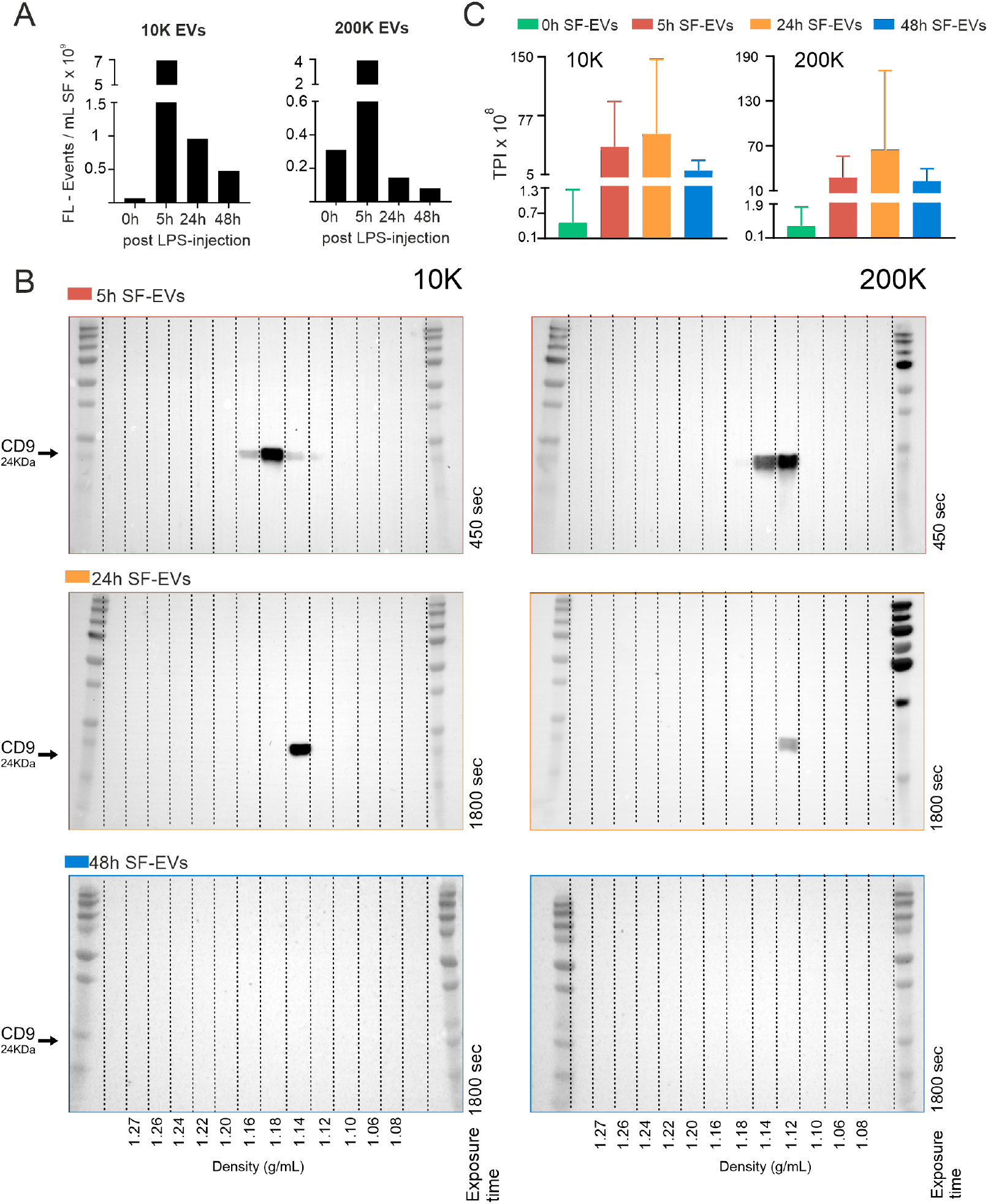
SF-EV characterization of donor samples used for lipidomics analysis. Synovial fluid was harvested at 5 hours, 24 hours, and 48 hours after an LPS insult. Synovial fluid-derived EVs were obtained at 10K and 200K, followed by a sucrose gradient. **A)** Sum of single EV-based high-resolution FCM of pre- & post-LPS injection (0h, 5h, 24h, and 48h) SF-EVs. Fluorescent events concentration in SF was calculated as the sum of single PKH67+ events measurements in sucrose gradient for the EV-enriched densities (1.10 to 1.20 g/mL). FL – Events: Fluorescent Events. **B)** Detection of CD9+ EVs by western blotting. Sucrose fractions of 10k and 200k-derived EV isolations were finally centrifuged at 200K to pellet EVs. Lanes were loaded with individually pelleted sucrose fractions. Blots were probed for CD9 (EV marker) and exposed for varying times, up to 30 minutes, to obtain a signal, even if weak. The upper blots represent a synovitis SF-EV sample at 5h (Red), the middle blots a 24h EV sample (Orange), and the lower a 48h EV sample (Blue). No signal was detected for either of the 48h SF-EVs blots. **C)** Lipid distribution from the EV-containing pooled sucrose densities 1.10-1.18 g/mL of three individual experimental horses. On the y-axis, the total phospholipid (relative) intensity (TPI) x 108 (mean +/- SD) is displayed. Synovial fluid was collected at 0 hours (pre-LPS)(Green), 5 hours (post-LPS) (Red), 24 hours (Orange), and 48 hours (Blue). Samples were analyzed in two lipidomics experiments (Experiment one: Horse 1: samples at 0h, 5h, 24h; Horse 2: 0h, 5h, 24h, experiment two: Horse 1: 48h; Horse 2: 48h; Horse 3: 0h, 5h, 24h, 48h). Larger standard deviations for 0h, 5h, and 24h SF-EVs are caused by inter-assay variations.

**Figure 9: Suppl. Fig. 4.**
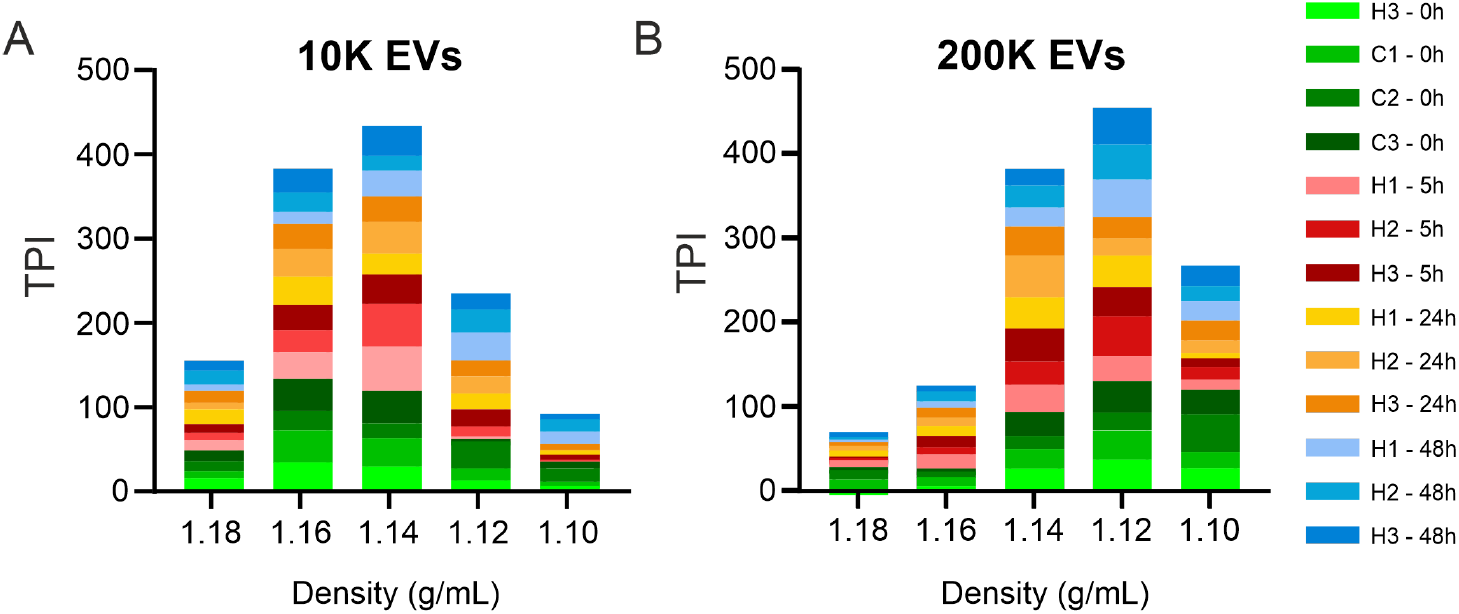
Normalized lipid intensities of EV samples shown in Supplementary Fig. 2C with additional healthy-SF EV samples. For each time point and EV-density fraction, samples were normalized by expressing the lipid intensity as a fraction of the sum of lipid intensities to define the lipid peak fractions of each EV sample. Zero hours SF-EVs in green tones (n=4, of which 3 were from healthy controls to compensate for the baseline levels below the detection limit in horses 1 and 2). Five hours SF-EVs are in tones of red, twenty-four hours SF-EVs in tones of orange, and forty-eight hours SF-EVs in tones of blue (n=3, H1, H2, and H3). **A)** Stacked bars plot of EVs pelleted at 10,000xg. Most EVs floated at densities of 1.16-1.14 g/mL. **B)** Stacked bars plot of EVs pelleted at 200,000xg. Most EVs peaked at densities of 1.14-1.12 g/mL.

**Figure 10: Suppl. Fig. 5.**
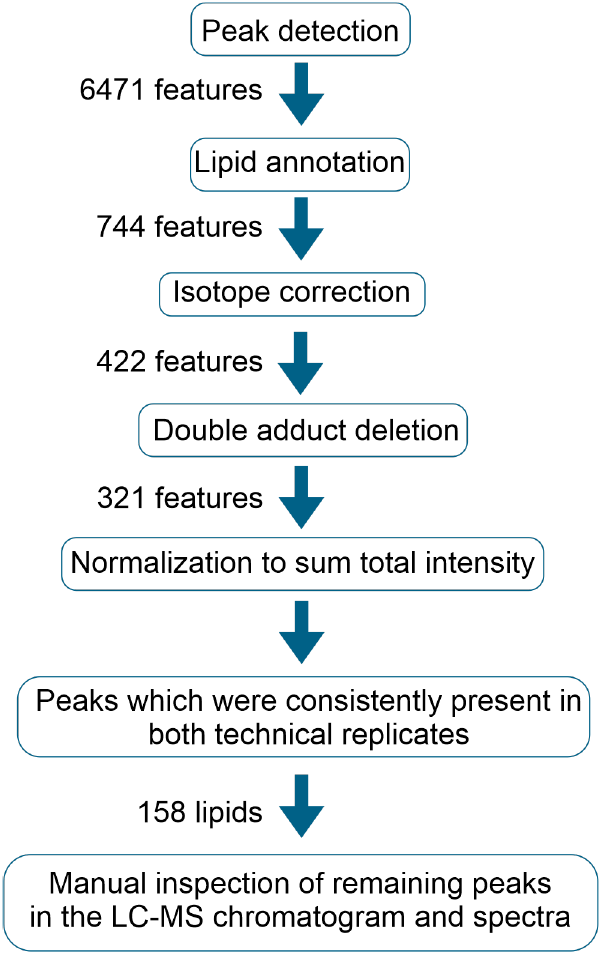
Workflow for lipid identification. Lipids were extracted from SF-EVs pelleted with successive ultracentrifugation steps of 10,000xg (10K) and 200,000xg (200K), followed by EV flotation in sucrose density gradients. In total EV from n=6 healthy SF samples (I.e.; n=3 pre-LPS-0h samples and n=3 additional healthy control SF samples to create robust baseline levels), n=4 synovitis SF samples post-LPS-5h (n=4; of which 2 were technical replicates between experiments to validate constant peaks), n=3 synovitis-resolution SF samples post-LPS-24h, and n=3 resolution SF samples post-LPS-48h were analyzed. Each n donor consisted of five fractions (F6-F10; densities 1.18-1.10 g/mL). Following MS analysis, the data were analyzed with the XCMS package from R, finding 6472 features, which were processed with a lipid annotation R script achieving the unmasking of 744 possible lipids. Lipid annotation is determined by the chromatographic mobility, the m/z ratio, and, when possible, the MS2 spectra. The features were corrected for isotopes; thus, 422 features were followed by double adduct deletion, reaching 321 features. The data were normalized by expressing all lipid species as fractions of the sum of the total lipid intensity of each biological sample (composed of 5 fractions). Furthermore, features were selected according to the consistency within the two technical replicates arriving at a total of 158 lipid species. The chosen final lipid peaks were inspected manually in the LC-MS chromatogram and MS2 spectra for confirmation of the species. Lastly, normalization within each density fraction was done (Suppl. Fig. 4) for selection of the samples for the phospholipidome analysis.

**Figure 11: Suppl. Fig. 6.**
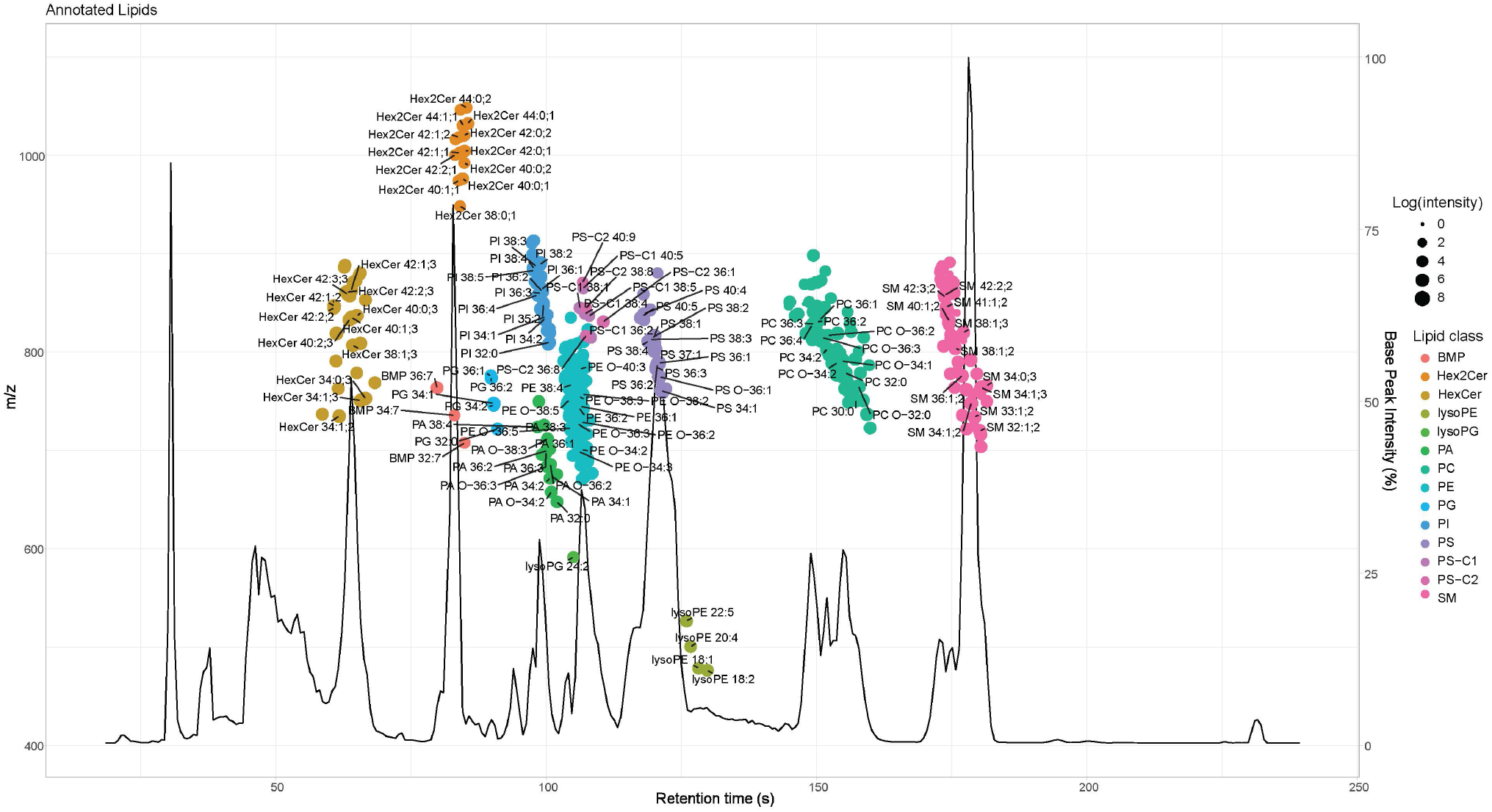
Base peak chromatogram with dot plot of the 422 features detected from synovial fluid-derived extracellular vesicles. The base peak chromatogram represents the mean chromatogram for all the samples, as described in Supplementary Fig 4, and analyzed in the mass spectrometer. The peak height illustrates the intensity as a percentage of the maximal peak (right y-axis). The dot plot depicts specific lipid species and their location according to their retention time (x-axis) and mass-to-charge ratio (left y-axis). Each color variation in the dot plot represents a lipid class. Some examples of potential annotation of lipids species are labeled with their possible respective adduct.

**Figure 12: Suppl. Fig. 7.**
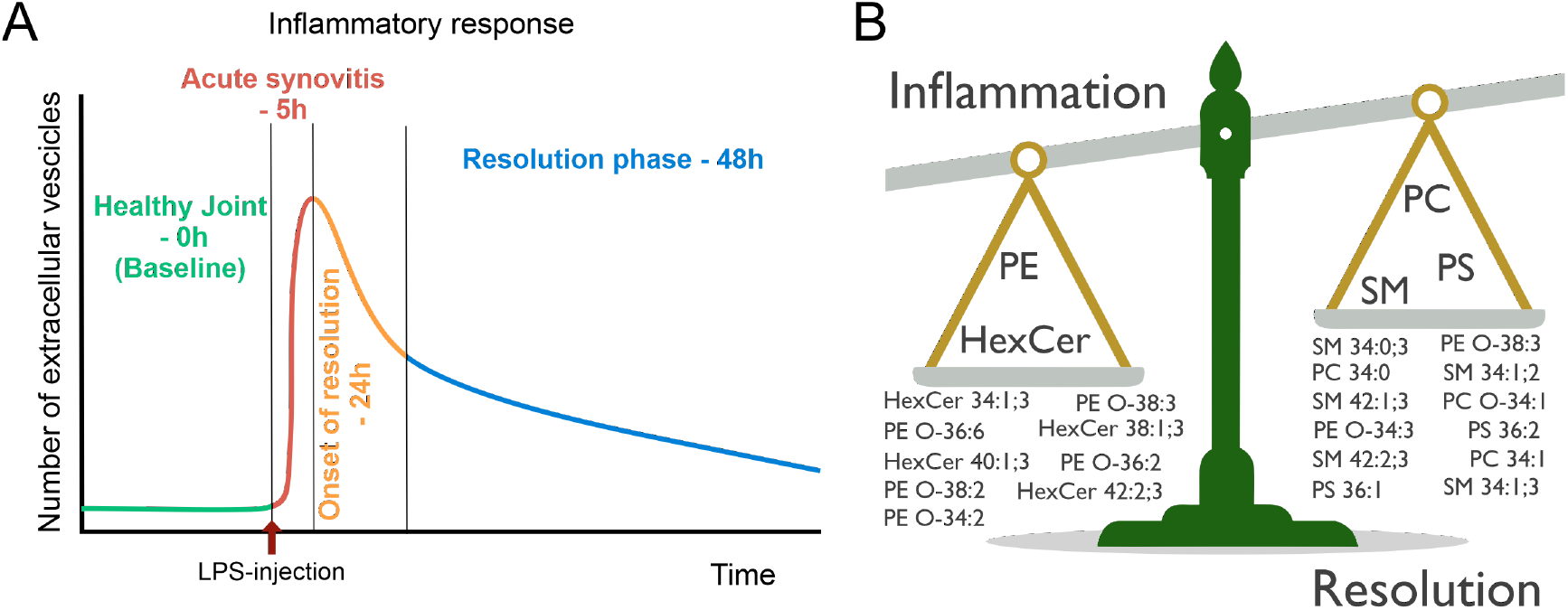
Schematic graphical representation of EV dynamics and balance in key lipids occurring during acute joint inflammation and its resolution. **A)** Representative cartoon of the EV dynamics during an inflammatory response. Starting with low EV levels in the healthy baseline situation (green), EV numbers peak (red) after the inflammatory stimulus (LPS injection; red arrow), followed by the onset of resolution and initial decrease of EVs (orange) and further EV reduction (blue) in the resolution phase. **B)** Illustrative lipid profile balance of important lipids during the inflammatory process. On the left of the balance, lipid classes and specific lipid species that emerge as dominant during the inflammation stages. On the right side of the balance are prevailing lipid classes and specific species that resurface in the resolution phase. Abbreviations: HexCer (hexosylceramide); PC, (phosphatidylcholine); PE, (phosphatidylethanolamine); PS, (phosphatidylserine); SM, (sphingomyelin).

